# Vitamin interdependencies predicted by metagenomics-informed network analyses validated in microbial community microcosms

**DOI:** 10.1101/2023.01.27.524772

**Authors:** Tomas Hessler, Robert J. Huddy, Rohan Sachdeva, Shufei Lei, Susan T.L. Harrison, Spencer Diamond, Jillian F. Banfield

**Affiliations:** The Innovative Genomics Institute at the University of California, Berkeley, California, USA; The Department of Earth and Planetary Science, University of California, Berkeley, California, USA; Environmental Genomics and Systems Biology Division, Lawrence Berkeley National Laboratory, Berkeley, CA, USA; The Center for Bioprocess Engineering Research, University of Cape Town, South Africa; The Future Water Institute, University of Cape Town, South Africa; The Department of Environmental Science, Policy and Management, University of California, Berkeley, California, USA

**Keywords:** Network analysis, metagenomics, microbial interactions, microbial ecology, *Variovorax*, vitamin exchange, thiamine

## Abstract

Microbial communities are fundamental components of most ecosystems but little is known about the interactions that structure them. Metagenomic data have revolutionized our understanding of complex consortia, yet predicted interactions are rarely explored experimentally. We developed an organism abundance correlation network to investigate factors that control community organization in soil-derived laboratory microbial consortia grown under dozens of conditions. The network was overlaid with metagenomic information about functional capacities to generate testable hypotheses. We developed a metric to predict the importance of each node within its local network environments relative to correlated vitamin auxotrophs and predicted that a *Variovorax* species is a hub because it is a highly important source of thiamine. Quantification of thiamine during the growth of *Variovorax* in minimal media conditions showed extraordinarily high levels of production of thiamine, up to 100 mg/L. This over-production of thiamine explains why the auxotrophs correlated with *Variovorax* are reliant on this organism for thiamine, despite the presence of other predicted thiamine producers in the community. A few of these thiamine auxotrophs are predicted to produce pantothenate that we show that *Variovorax* requires to grow, implying that a subset of vitamin-dependent interactions are mutualistic. We found that *Cryptococcus* produces the B-vitamin pantothenate, and co-cultures led to a 90-130-fold fitness increase for both *Cryptococcus* and *Variovorax*. This study demonstrates the predictive power of metagenome-informed, microbial consortia-based network analyses for identifying microbial interactions that underpin the structure and functioning of microbial communities.

## Introduction

Microbial communities are critical to human health, agriculture, industrial bioprocesses and the functioning of natural environments such as soil, groundwater and oceans. However, our ability to predict the structures of microbial communities, and how these might be altered by perturbations such as sustained environmental changes, is lacking. Microbial communities are underpinned by numerous interactions, the understanding of which should enhance our ability to manipulate and improve microbiome functioning.

Correlation-based network analysis is one of the longest-standing methods for identifying interacting organisms in ecosystems (Proulx et al., 2005). In the context of microbial communities, such networks and interactions have, by and large, been predicted using abundance and taxonomic information inferred from 16S rRNA gene amplicon sequencing data (as reviewed by Matchado et al., 2021; Banerjee et al., 2018). However, 16S rRNA sequencing does not provide any information about an organism’s functional potential and also suffers from the fact that the network nodes representing the quantifiable units of 16S analysis (amplicon sequence variants or operational taxonomic units) are not necessarily synonymous with independent microbial species. Thus, networks constructed from 16S data likely do not represent co-occurrences of unique and independent species as well as lack any functional basis to explain why some organisms are network hubs or exhibit strong interdependencies with other organisms.

To generate a mechanistic understanding of interactions in communities, Qian et al. (2021) recently suggested moving away from correlation-based networks to modeling approaches. These include Generalized Lotka-Voltrerra models (Connors et al., 2022) and Dynamic Flux balance analysis models (Dukovski et al., 2021) that are based on genomic information. For example, genomic information has been used to generate metabolic models that predict the interactions between two species in co-culture (Freilich et al., 2011) and have demonstrated that cross-feeding is common. The inherent nature of cross-feeding between organisms that do not encode for the biosynthesis of their own metabolites necessary for growth gave rise to the Black Queen hypothesis (Morris et al., 2012). This hypothesis supposes that the loss of genes and pathways enables the conservation of an organism’s resources and leads to microbial dependencies.

Despite the progress and the promising outlook of these modelling approaches, they remain difficult to implement for complex microbial communities under varied conditions, and are therefore, largely limited to simple microbial systems. In contrast, we hypothesized that the functional basis for microbial interactions can be predicted if genome-resolved metagenomic data for complex microbial communities are incorporated into conventional network analyses. In this study, we constructed a correlation-based network for bioreactor-grown microbial communities using previously published metagenome datasets (Kantor et al., 2015, Kantor et al., 2017, Rahman et al., 2017, Huddy et al., 2021) and overlaid functional information onto this network. The bioreactors were inoculated with thiocyanate-degrading (SCN^-^) microbial culture originating from a mine tailings reservoir. As vitamin B12 (cobalamin) is produced by only a subset of organisms within these systems (Huddy et al., 2021), we hypothesized that vitamin production may underlie the functioning of specific organisms as hubs in this network. Vitamins are essential for cellular function, and vitamin auxotrophs are frequently observed within microbial communities (Rodionov et al., 2003; Sokolovskaya et al., 2020). Thus, we focussed on vitamin cross-feeding. Sokolovskaya et al. (2020) proposed cobamides as a model system for studying microbial interactions, in part because their biosynthetic pathways are well annotated. Experimental evidence supports this focus. For example, a prior study generated vitamin, amino acid, nucleotide and carbon metabolism auxotrophs in *E. coli* and found that vitamin auxotrophs were more likely to enter into cross-feeding interactions with organisms with complementary auxotrophies in mixed communities (Guillen et al., 2021). Using our approach we identified organisms that are hubs in the correlation network, including one that is predicted to supply thiamine to correlated thiamine auxotrophs. We tested this prediction by isolating this predicted thiamine producer and thiamine auxotrophs and by performing co-culture experiments in which the amount of thiamine produced was quantified. We suggest that the overlay of genome-resolved metagenomic functional information onto network analyses can more generally explain the metabolic interactions that occur within subnetworks of microbial communities.

## Results

### Microbial community network structure

This study leveraged time series, genome-resolved metagenomic datasets that were acquired from bioreactors that were inoculated with a consortium established primarily from soil and operated over a range of conditions, including varying thiocyanate (SCN) concentrations, applied hydraulic retention times, and with or without the addition of molasses as an additional energy source (Kantor et al., 2015; 2017; Rahman et al., 2017; Huddy et al., 2021). Because the bioreactors contained organisms able to degrade thiocyanate, the thiocyanate concentrations remained below detection levels, preventing any toxicity effects. The 92 metagenomic datasets included draft genomes for virtually all of the relatively abundant organisms. We used this data to construct an abundance-based co-occurrence network onto which we overlaid functional capacity predictions, predicted vitamin auxotrophies, and identified and tested these predicted microbial interactions (Figure 1).

**Figure 1.**
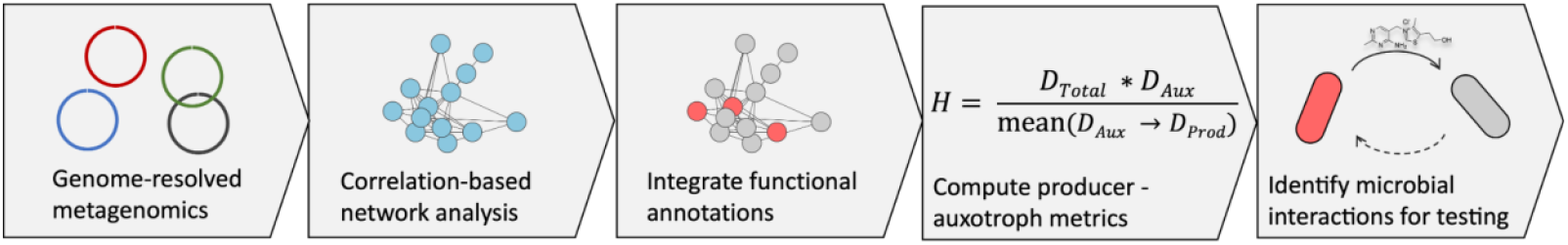
Schematic diagram of the computational and statistical approach employed to identify microbial interactions using genome-resolved metagenomics and network analyses in this study. First, metagenome-associated genomes are recovered and used for abundance-based correlation network analysis (panels 1 and 2). Functional traits, such as vitamin biosynthesis, are inferred from the genomes and overlaid onto the network (panel 3). The importance of nodes with a given functional annotation is estimated (panel 4). The generated metric estimates the importance of a vitamin-producing organism for the provisioning of this vitamin to other members in the local network neighborhood. This leads to the generation of hypotheses describing potential microbial interactions (panel 5).

We used positive linear correlations between the 309 species-representative genomes recovered across the 92 bioreactor metagenomes (Figure 2A, see methodology). Organisms which had strong positive correlations were included in the correlation network. This network has 120 nodes that represent independent microbial species, with an average of four neighbors per node. The total network density was just 6.7% (the percentage of total possible edges between the nodes in a network), indicating that the network was sparse (low level of connectivity), with a modularity of 0.52 (a measure of the separation of nodes within a network into interconnected groups), which is due to the extensive separation of groups of organisms into sub-networks. We delineated eight nodes as ‘network hubs’ (Figure S1) based on elevated betweenness centrality, a whole-network metric that represents the relative importance of a node for the overall connectivity within a network. The hubs were three Alphaproteobacteria (a *Mesorhizobium sp*., a *Sphingomondales sp*., and *Rhodospirillales sp*.), three Betaproteobacteria (*Variovorax, Commamonas, Nitrospira*), a member of the Gammaproteobacteria family Rhodanobaceraceae and a member of the Bacteroidiota family Chitinophagaceae.

**Figure 2.**
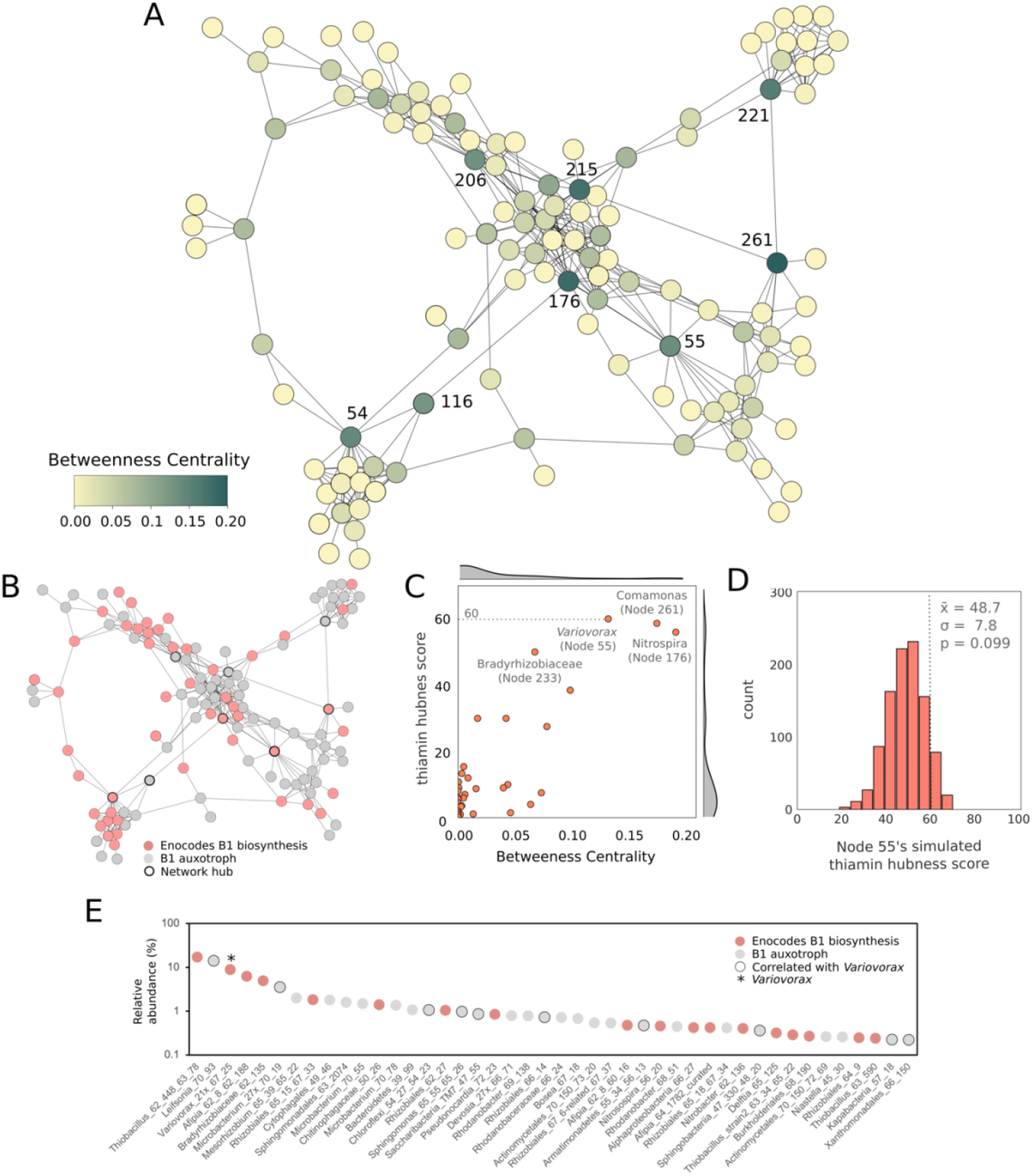
*Variovorax* (node 55) is a network hub predicted to be important for thiamine production in the bioreactor communities. (A) Network analysis based on sparCC correlation across 92 metagenomes was used to predict associations of microorganisms with other members. The nodes with the greatest betweenness centrality were delineated as “network hubs” and are numbered. Four of these network hubs (B) encode genes for the biosynthesis of thiamine. Of these four hubs (C) *Variovorax* (Node) 55 shows the greatest importance for correlated auxotrophic members based on “thiamine-hubness metrics” which considers the degree, connections to B1-deficient nodes, and the number of connections those B1-deficient nodes have to B1-producers. (D) Distribution of calculated vitamin hub metric scores of Variovorax (Node 55), given the network shown in Figure 2A, after randomly assigning 38 nodes within the network the capacity to produce thiamine, over 1000 simulations. (E) *Variovorax* is present in communities with many other bacteria able to produce thiamine but is correlated almost exclusively with thiamine auxotrophs within bioreactor communities. The rank abundance curve of the relative abundance of the 44 most prevalent microorganisms in bioreactor sample SCN18_30_10_14_R1_P is shown as an example of this phenomenon.

The network was partitioned into modules, subnetworks which make up the network, by computing the maximum modularity of the network based on Louvain heuristics (Figure S1). Modular networks such as this have a greater number of edges within these modules than they do with other modules. Four of the modules have a single hub organism, two lack a hub organism and two contain two hub organisms.

We quantified the possible importance of vitamin producers within their local network neighborhood by defining a node-level “vitamin-hubness” metric. The metric computes the product of a node’s degree (number of connections) and its number of connections with vitamin auxotrophs, relative to the mean number of connections these auxotrophs have with vitamin producers. This identifies well-connected vitamin producers which are connected to many auxotrophs. The second term reduces the score of producers whose auxotrophs are connected to other producers from which they could obtain the vitamin of interest. Using this approach we identified multiple organisms as potentially important for the provisioning of B vitamins. These included the hub organisms *Nitrospira* (Node 176), *Comamonas* (Node 261) and *Variovorax* (Node 55) for the provisioning of thiamine, *Chitinophagacaea* (Node 221) for pantothenate, as well as a *Rhodanobacter* (Node 48) for the possible provisioning of biotin and riboflavin (Figure S2, S4).

We performed an isolation campaign to recover as many organisms represented in the network as possible. This led to the isolation of 43 bacterial species and single yeast species. Only one hub organism was isolated, a *Variovorax* corresponding to Node 55. The genome sequence of the isolate corresponded (99.99% ANI) with the genome from the metagenome for the hub organism. We, therefore, focused on the subnetwork for which *Variovorax* is the hub.

*Variovorax* was present in communities with many other predicted thiamine producers, but the majority of the organisms which were correlated with *Variovorax* were thiamine auxotrophs (Figure 2E). Interestingly, each module contained at least one metagenome-assembled genome (MAG) with a complete pathway for the biosynthesis of thiamine; most were Alphaproteobacteria or Betaproteobacteria, (Table S1, Table S2, Figure S3). Many members of the communities where *Variovorax* was present were also predicted to produce thiamine (Figure 2E). This raises the question of why *Varivorax* rather than another thiamine producer serves as the hub organism.

We wanted to determine if the reason for *Variovorax’s* high “thiamine hubness” of 60.0 was due to the enrichment of thiamine auxotrophs in its local network neighborhood, or due to the absence of other thiamine producers aside from *Variovorax*, or a combination of both. We randomly assigned 37 nodes, as well as Variovorax, as thiamine producers (38 is the true number of nodes found to be capable of thiamine production) and 82 as thiamine auxotrophs and recalculated *Variovorax’s* thiamine-hubness score. We repeated this process for a total of one thousand simulations (Figure 2D). The distribution of these scores resembles a normal distribution with an average score of 48.8 and a standard deviation of 7.8. *Variovorax’s* thiamine-hubness score of 60.0 is elevated with the respect to the mean but was found not to be significantly greater. Therefore, *Variovorax* is partially enriched with nodes auxotrophic for thiamine, and these auxotrophic nodes are connected to few thiamine producers other than *Variovorax*.

### *Variovorax* thiamine biosynthetic operon

A single thiamine-pyrophosphate (TPP) riboswitch-regulated operon contains the majority of the thiamine biosynthetic genes within the *Variovorax* genome. Included in this operon are a thiamine permease and a TonB-dependent receptor-like protein predicted to be a thiamine active transporter (Figure 3). The biosynthetic genes include the *thiD* whose product is responsible for phosphorylation of the pyrimidine moiety, several genes involved in the biosynthesis of the thiazole ring and *thiE* responsible for the linking of these two moieties. The *thiC* gene encoding for the phosphomethylpyrimidine synthase, responsible for the generation of the hydroxymethyl pyrimidine, is found in a separate TPP riboswitch-regulated operon. The remainder of the genes involved in thiamine biosynthesis (genes *iscS*, *thiL*, *dxs*) are located separately in the *Variovorax* genome. Neither the standard thiamine transporter genes *thiT* (also referred to as *yuaJ*) nor any of the *ykoFEDC* genes were identified in the *Variovorax* genome, possibly due to poor transporter annotation.

**Figure 3.**
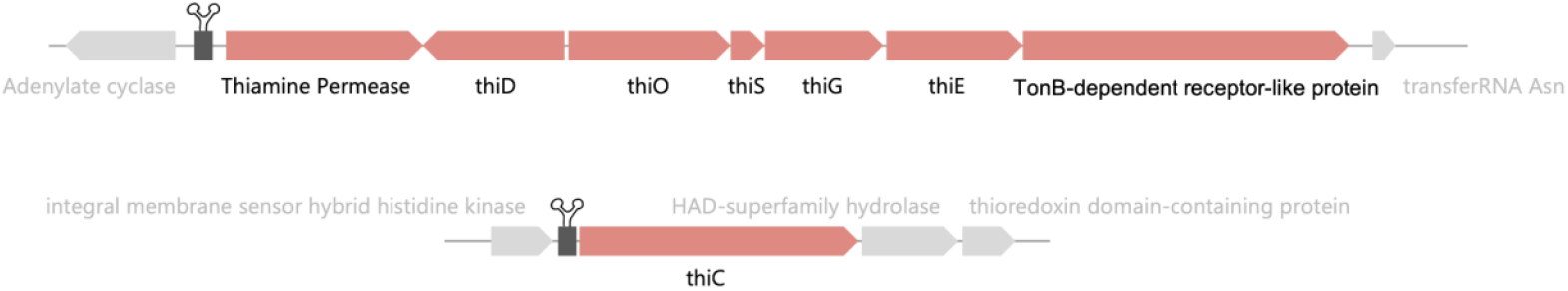
Schematic diagram of the thiamine biosynthesis operons regulated by TPP riboswitches in the isolated *Variovorax* genome.

Dysregulated thiamine operon mutants have been generated for biotechnological application (Schyns et al., 2005). This requires the mutation of three genes, one of which is the thiamine-monophosphate kinase (*thiL*). The isolated *Variovorax thiL* amino acid sequence appears functional, based on a comparison with a functional *thiL* sequence analysis performed by McCulloch et al. (2008). The active site residues of our Variovorax’s *thiL* are present and the additional conserved residues identified by McCulloch and co-authors were largely maintained (Supplementary materials).

### The *Variovorax* genome encodes several capacities potentially important for interactions

The 7.9 Mbp *Variovorax* genome is predicted to encode for a highly versatile aerobic metabolism, with the capacity to utilize a variety of sugars (including trehalose and starch), lipids, and complex and recalcitrant (often plant-derived) carbon compounds such as xylan and cellulose (Figure 4). The genome also encodes for the biosynthesis of thiamine, the siderophore Serobactin, fatty acids, as well as encoding for a large number of genes relating to motility and chemotaxis. However, the genome lacks a complete pathway for the biosynthesis of pantothenate (vitamin B5). The *Variovorax* isolate grew aerobically on sugars, pyruvate and branched-chain amino acids, but growth required the addition of pantothenate acid (vitamin B5). *Variovorax* was found to be correlated with two pantothenate acid-producing bacteria, a *Sphingobacteria* and a *Kapabacteria*.

**Figure 4.**
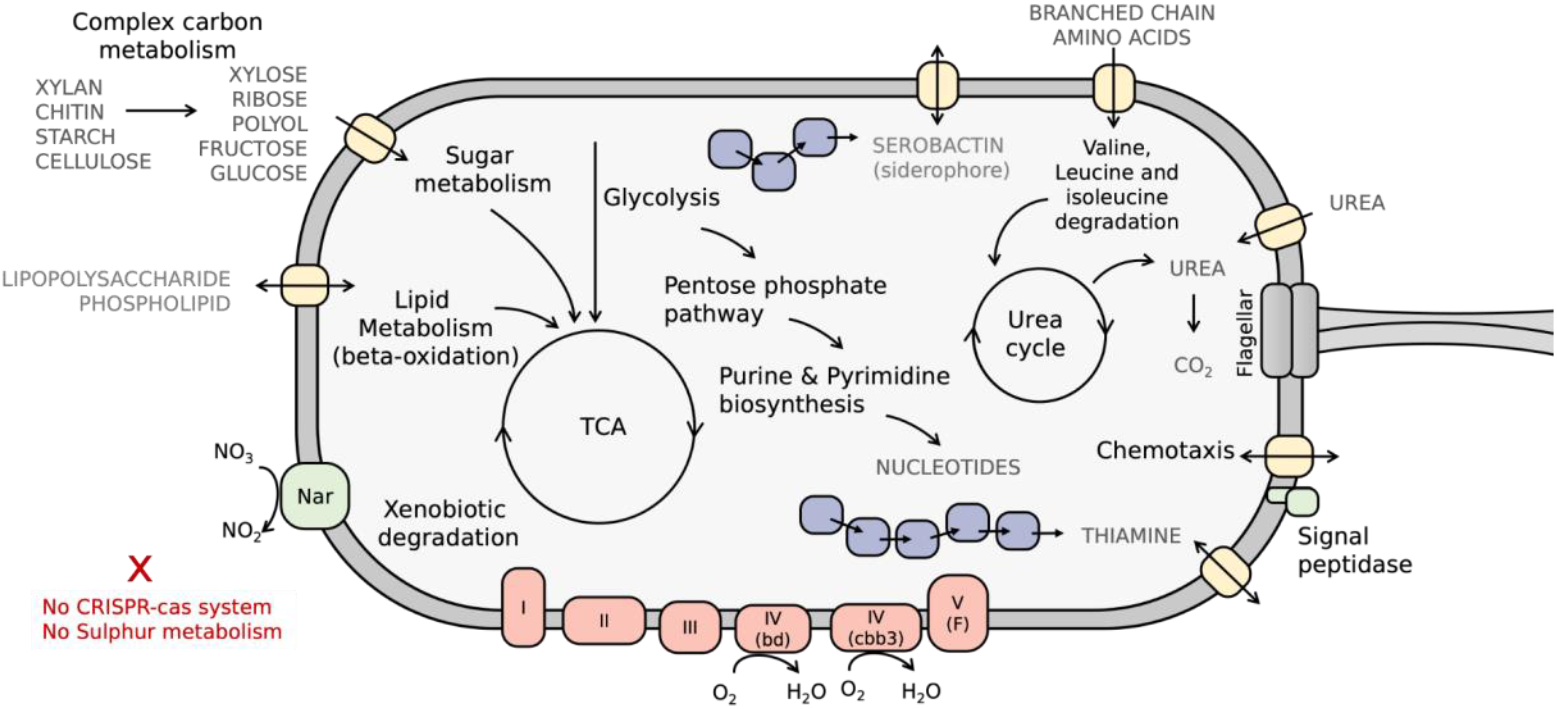
Cell diagram representing the genetic potential of the *Variovorax* isolated from the bioreactor community. *Variovorax* is predicted to be metabolically versatile, can produce the siderophore Serobactin and has many genes involved in chemotaxis. These encoded genes and pathways provide many possible mechanisms for interactions with other members.

### *Variovorax* supports *Saccharibacteria* and other members of enigmatic phyla

A number of bacteria correlated with *Variovorax* belong to rare and understudied phyla including a *Chloroflexota*, a *Kapabacteria*, an *Armaimondota* and two *Saccharibacteria* (Figure 5A). As is true with other *Saccharibacteria*, the *Saccharibacteria* have small genomes and highly reduced metabolic capacities. The host of the *Saccharibacteria* (Node 45) was previously predicted to be a *Microbacterium ginsengsoli* (Node 144) based on co-occurrence patterns identified through hierarchical clustering (Huddy et al., 2021). Neither *Saccharibacteria* nor *Microbacterium* have genomes that encode for the biosynthesis of thiamine, but nonetheless, all encode several genes coding for proteins which are known to require thiamine as a cofactor. The proliferation of *Saccharibacteria* to high abundance levels only occurred when *Variovorax* was reasonably abundant. This suggests that *Saccharibacteria* are dependent on both the host *Microbacterium* and *Variovorax*.

**Figure 5.**
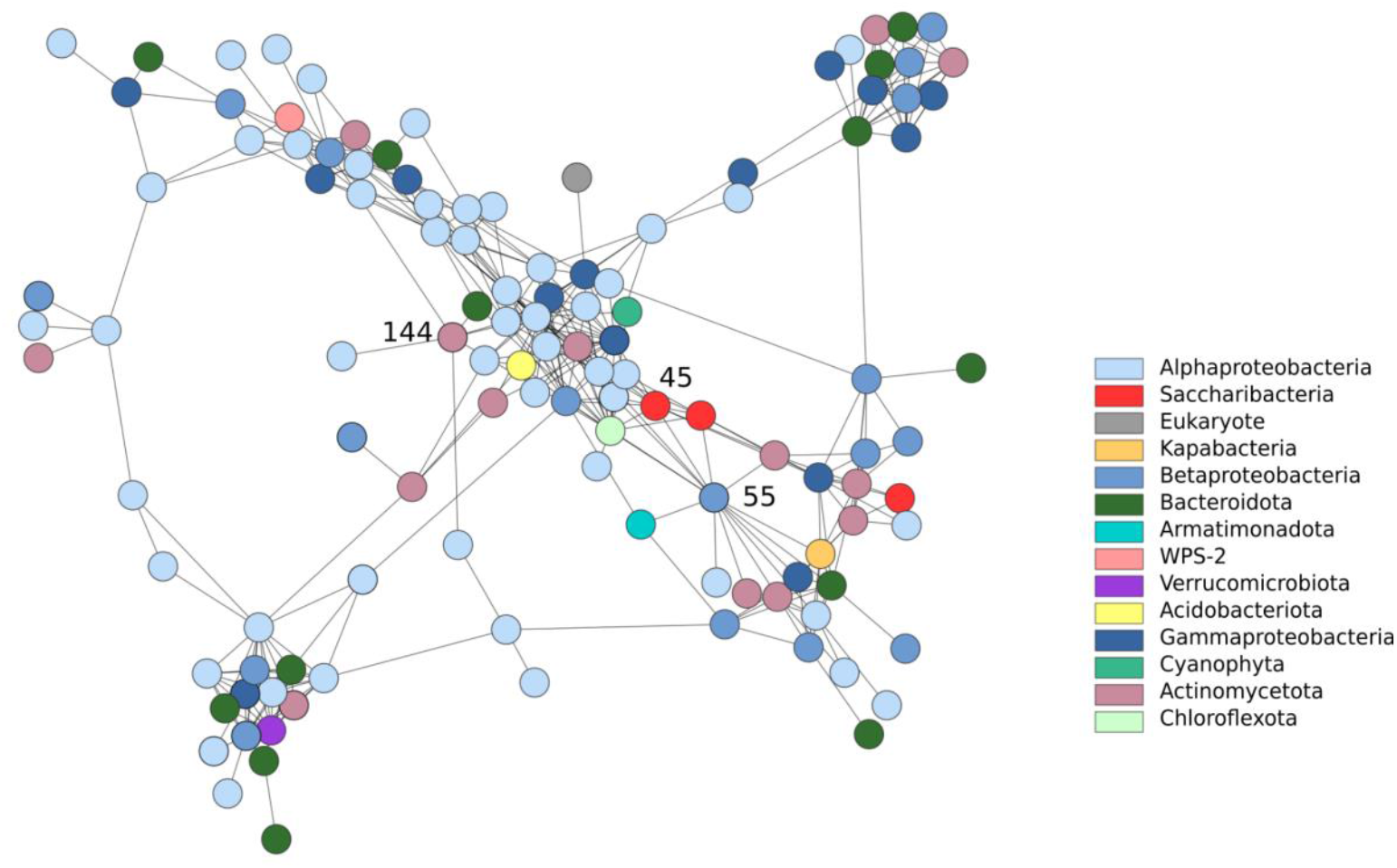
*Variovorax* (Node 55) is correlated with a *Saccharibacteria* (Node 45) as illustrated in the network plot showing phyla annotations. *Microbacterium* (Node 144) and *Saccharibacteria* both have multiple genes which encode proteins which require thiamine as a cofactor - neither of these genomes encodes for thiamine biosynthesis. A version of this figure with clearer node separation can be found in the supplementary material (Figure S5).

### Thiamine biosynthesis is common in the genus *Variovorax*

*Variovorax* has been suggested as an important hub in rhizosphere communities due to its capacity to degrade complex carbon compounds, but the findings of this study raise the possibility that vitamin production may also contribute to its hub behavior. Thus, we investigated whether thiamine production is a common trait within the *Variovorax* genus. We assessed all *Proteobacterial* genomes in GTDB for their capacity to produce thiamine. The capacity to synthesise thiamine was observed in 24.5% of Gammaproteobacteria genera, 20% of Proteobacterial genera and only 10% of all 9884 Proteobacteria genomes.

Of the 72 isolate and metagenome-derived *Variovorax* genomes in GTDB, 27 encoded a near-complete to complete thiamine biosynthetic pathway (37.5 %). Each of these 27 genomes encoded a thiazole synthase (*thiG*), 17 of which encode a complete thiamine biosynthesis pathway and the other 10 encode a near-complete pathway (Figure 6), as defined by the presence of genes in four major enzymatic steps (described in the methodology). In contrast, only five were found to encode a complete pantothenate biosynthesis pathway.

**Figure 6.**
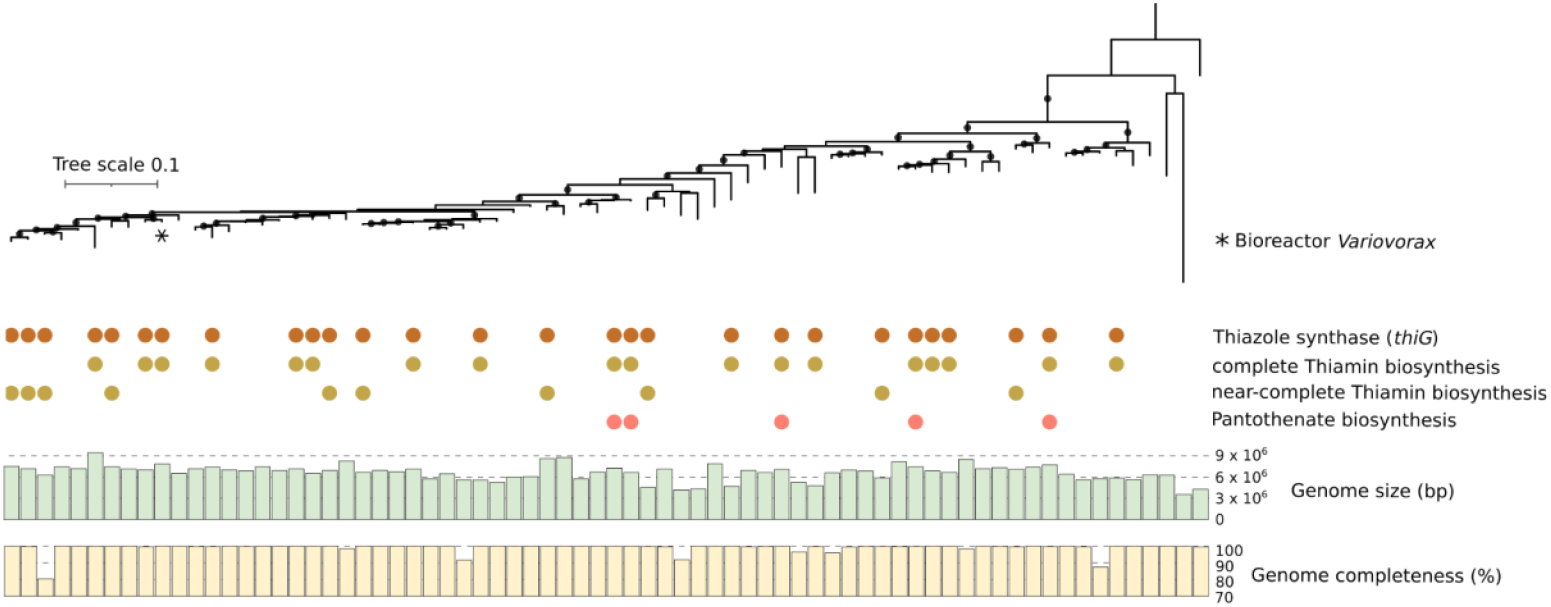
Thiamine biosynthesis is common in *Variovorax* genomes. A phylogenetic tree of 16 concatenated ribosomal proteins from 70 *Variovorax* genomes deposited in GTDB together with the *Variovorax* isolated from our bioreactor. The encoding of a complete Vitamin B1 biosynthetic pathway is shown and was found to be encoded in 27 of the 72 GTDB *Variovorax* genomes. Only five of the genomes encode a complete pantothenate biosynthetic pathway. *Rhodoferax saidenbachensis* is shown as the outgroup. Branches supported by bootstrap values greater than 75 are shown as circles. The bioreactor *Variovorax* shares the greatest similarity with *Variovorax sp003019815* (GCF_003019815.1).

### *Variovorax* increases the fitness of thiamine auxotrophs through thiamine production

To test the hypothesis that *Variovorax* can act as a thiamine supplier to thiamine auxotrophs we screened our bioreactor isolates for microorganisms whose growth was greatly enhanced by the incorporation of thiamine into minimal media. This identified the yeast isolate belonging to the genus *Cryptococcus* as a thiamine auxotroph (Figure 7A). We hypothesized that in co-culture in minimal media, the predicted thiamine producer *Variovorax* would confer a fitness advantage to the *Cryptococcus*. Indeed, the cell density of *Cryptococcus* increased ~90-fold in a co-culture (Figure 7B). *Variovorax* too benefitted from this interaction, increasing from 7×10^5^ cells/ml in pure culture to 9×10^8^ cells/ml in co-culture with *Cryptococcus* (~130-fold).

**Figure 7.**
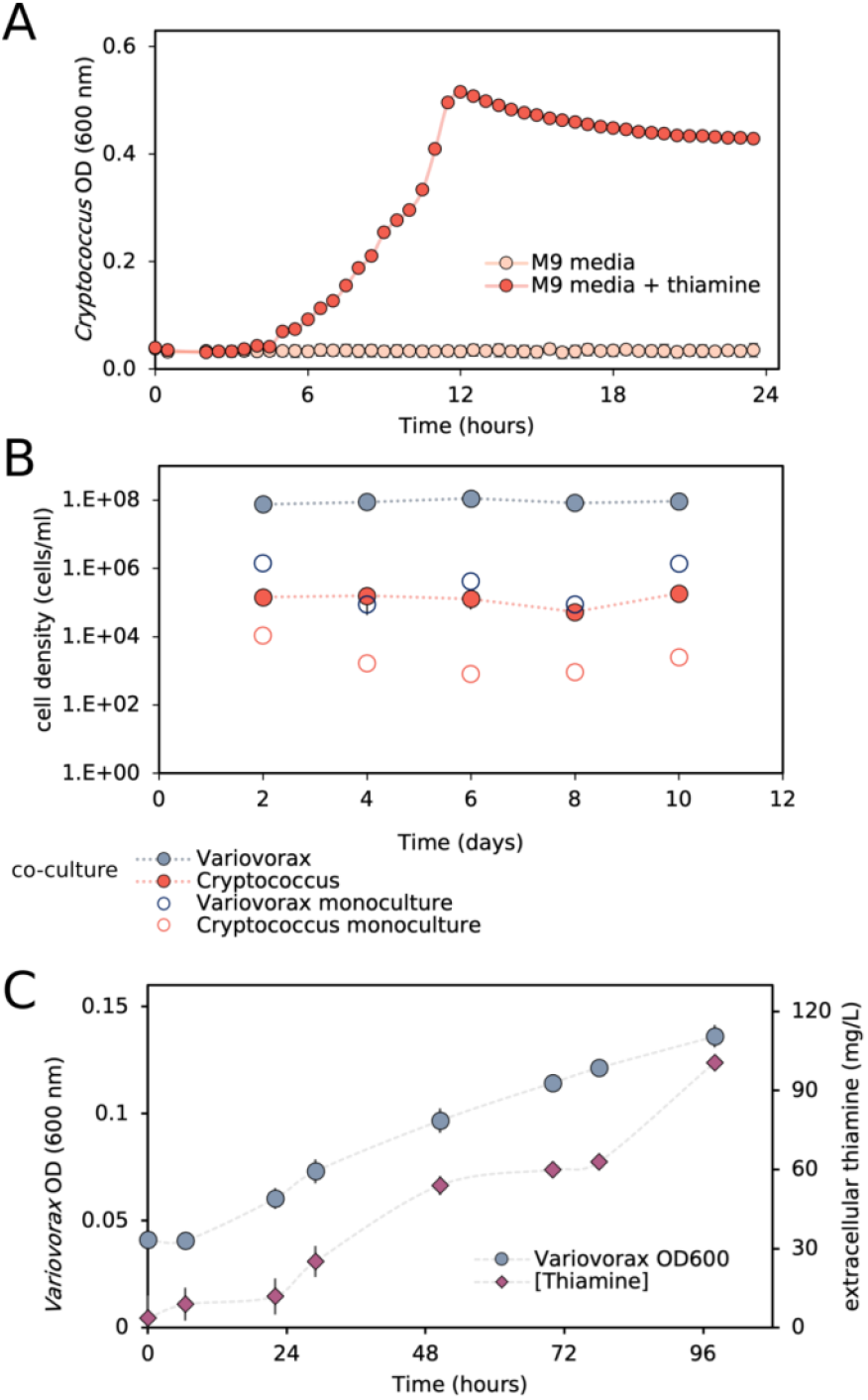
(A) A *Cryptococcus* isolated from the bioreactor community is shown to be a thiamine auxotroph through supplementation of thiamine M9 minimal media. (B) The steady-state cell density of this *Cryptococcus* increases ~50-fold when grown in co-culture with *Variovorax* in M9 media. *Variovorax* too receives a fitness advantage from the interaction. (C) *Variovorax* produces large quantities of extracellular thiamine in pure culture in minimal media supplemented with 25% spent media from *Cryptococcus* grown in minimal media for 5 days. Error bars represent one standard deviation from the mean of 3 replicates.

To confirm that Variovorax was producing thiamine we monitored for thiamine while culturing *Variovorax* in minimal media with the addition of 25% spent media from a *Cryptococcus* five-day monoculture in minimal media. In minimal media without the addition of spent media, no visible growth of *Variovorax* was observed. The spent media enabled *Variovorax* to reach an optical density (OD_600_) of ~0.15 after 96 hours (Figure 7C), but in an almost linear fashion. Thiamine concentration increased simultaneously with *Variovorax* growth from 3.6 mg/L at the start of the experiment to a maximum of 100 mg/L after 96 hours.

We previously predicted Variovorax to be a pantothenate auxotroph based on its genome and confirmed this in a series of short growth experiments (Table S3). Despite the poor growth of the *Cryptococcus* in minimal media over five days, we were able to detect ~5 mg/L of pantothenate in its spent media, suggesting that *Cryptococcus* production of pantothenate may account for *Varioovrax’s* improved growth in the co-culture.

### *Variovorax* produces thiamine during lag phase

We performed growth experiments of *Variovorax* in minimal media in the presence of increasing pantothenate concentrations (Figure 8). In all instances, slow linear growth was initially observed followed by exponential growth after an extended lag phase (Figure 8A). Increasing pantothenate concentrations allowed greater optical densities to be reached by the onset of this lag phase. Subsequently, extracellular thiamine production proceeded linearly (Figure 8B), until the start of exponential growth, upon which the produced thiamine was rapidly consumed. Higher concentrations of supplied pantothenate corresponded with greater produced thiamine concentrations, with a maximum of 39.5 mg/L thiamine observed in the 5 mg/L pantothenate supplemented cultured. The higher concentrations of pantothenate also allowed exponential growth to be reached sooner after inoculation. However, the specific thiamine production rate (thiamine produced per unit optical density) was largely consistent between supplied pantothenate conditions. Interestingly, *Variovorax* showed very little growth within the first 144 hours in the absence of supplied pantothenate, yet still produced considerable quantities of extracellular thiamine (8.4 mg/L) over the first 144 hours.

**Figure 8.**
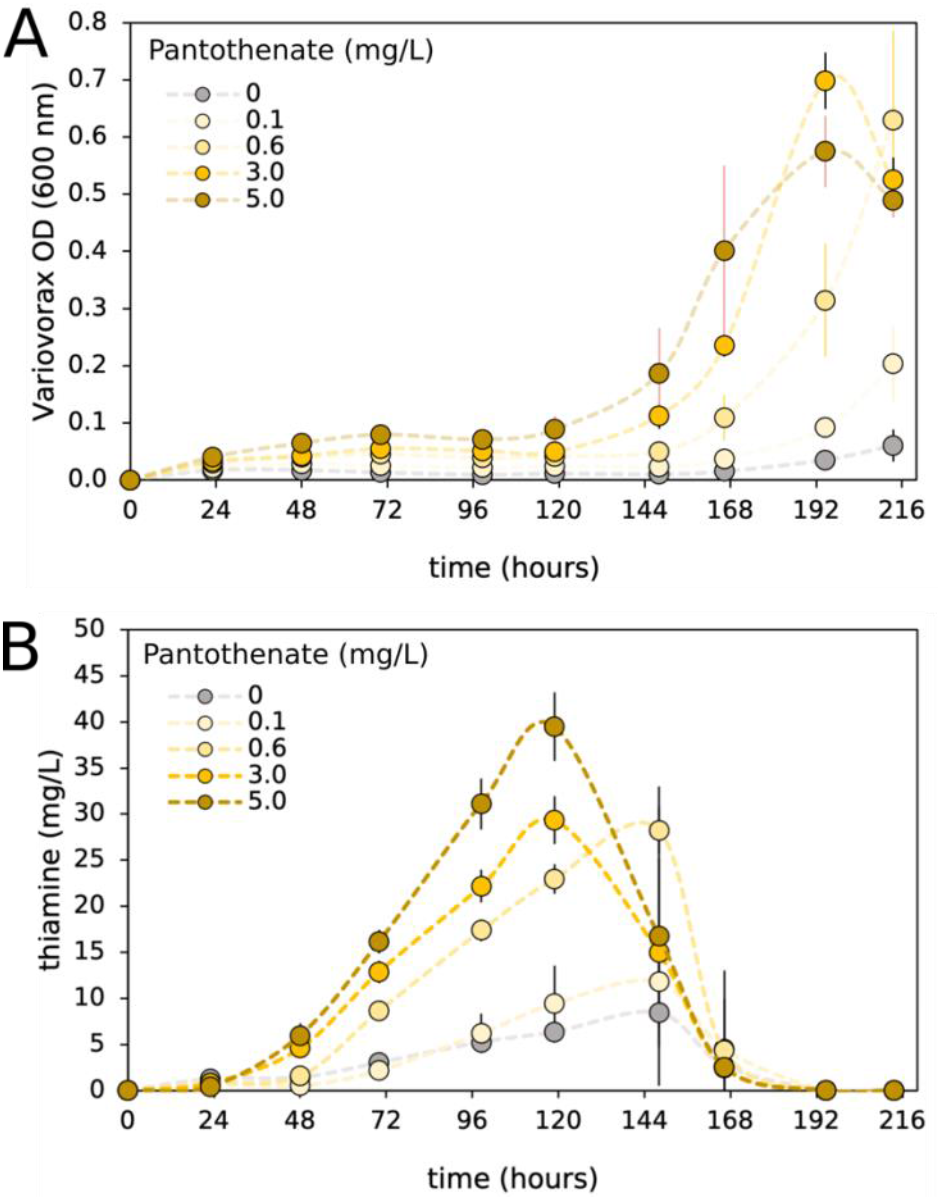
Growth of *Variovorax* (A) and its thiamine production (B) when grown in M9 minimal media with the addition of varying concentrations of pantothenate (vitamin B5). Error bars represent one standard deviation from the mean of 4 replicates.

## Discussion

Here by overlaying functional metagenome information onto co-occurrence networks we identified several microorganisms in a complex microbial community which we predict to be important for the provisioning of B vitamins. We developed a “hubness” metric that can be used to evaluate the extent to which a resource of interest is important for structuring microbial interactions within subnetworks of networks that describe microbial community co-occurrence patterns (Figure 2C). We used this approach to calculate the “thiamine hubness” metric and predicted that a *Variovorax* sp. is a hub (Figure 2A) because it supplies thiamine to other organisms in its immediate network vicinity. The metric considers the number of connections between a hub organism (such as *Variovorax*) and other organisms in its immediate network neighborhood, the number of thiamine autotrophs in this neighborhood set, but it also considers the number of connections between these thiamine auxotrophs and predicted thiamine producers other than *Variovorax*. This last term downplays the importance of a predicted interaction in cases where other organisms (within the immediate network neighborhood or outside of it) could be alternative sources of the resource. This metric can be readily applied to future analyses to assess auxotrophies and metabolic exchange in other microbial communities.

We isolated a network hub organism *Variovorax*, which we predicted to be important for the biosynthesis of thiamine. Thiamine is a cofactor required by enzymes throughout central metabolism including those of the citric acid cycle (Reed and Cox, 1966), pentose phosphate pathway (Racker et al., 1953), and amino acid catabolism and anabolism (Schloss, 1992), yet not all bacteria in the bioreactor communities are predicted to encode the genes necessary for thiamine synthesis. Unlike most prior studies that have used genomic data to investigate potential microbial interdependencies, we tested the prediction experimentally and correctly predicted thiamine production and interdependence. We confirmed vitamin exchange through isolation and characterisation of the *Variovorax* predicted to be responsible for thiamine production, together with an isolated thiamine auxotroph, *Cryptococcus. Cryptococcus* was not present within the correlation network, likely due to its occurrence in only a handful of communities. However, we report evidence that *Cryptococcus* is a pantothenic acid producer, a B-vitamin for which *Variovorax* is auxotrophic. This complementarity led to a 90 - 130-fold increase in cell density for both members when grown in co-culture.

A limitation of our approach is that we did not consider non-linear relationships when constructing the correlation network (Matchado et al., 2021). It is plausible that other possible interactions of *Variovorax* could be elucidated without reliance on linear regression alone. An example of this could be the provision of siderophores, which this genus is commonly known to produce (Kurth et al., 2016), but may not be detectable as linear relationships.

We noted a correlation between *Variovorax* and two *Saccharibacteria*, a *Kapabacteria*, a *Chloroflexi* and an *Arminatododetes sp*. (Figure 5). The predicted *Microbacterium* host of one of the *Saccharibacteria* is also predicted to be a thiamine auxotroph. These findings raise the important possibility that the commonly predicted dependence of *Saccharibacteria* on an

*Actinobacteria* species (Cross et al., 2019; Bor et al., 2020; Murugkar et al., 2020; Batinovic et al., 2021) can be more complex, in this case involving a third organism capable of vitamin production (*Variovorax*, which supplies thiamine).

*Variovorax* has been shown to be an important member of rhizosphere microbial communities (Finkel et al., 2020). Previously reported *Variovorax* genomes encode a repertoire of complex carbon compound degrading genes, which is consistent with the genetic potential of our *Variovorax* (Figure 3). Members of this genus have been shown experimentally to degrade complex carbon compounds such as herbicides (Dejonghe et al., 2003), and plant-produced isoprenes (Dawson et al., 2022), the breakdown products of which support other microbial members unable to directly consume these compounds. Our results are the first to demonstrate *Variovorax* supports microbial communities through the provision of a vitamin. We found the thiamin biosynthesis operon is present in many members of the genus *Variovorax* (Figure 6) and thus thiamine production may be a common function, in addition to its multiple other, already elucidated, roles.

*Variovorax* has been implicated in symbioses involving plants as well as bacteria. Finkel et al. (2020) found that *Variovorax* was the sole member of a 185-member syncom responsible for the degradation of the bacteria-produced auxin which, unmetabolized, leads to root growth inhibition in *Arabidopsis* seedlings. *Variovorax* has also been noted frequently as a good plant endosymbiont (Reiter et al., 2006). An endophytic *Herbaspirrilum* was similarly found to produce a serobactin siderophore found in our *Variovorax* genome and was shown to directly aid in plant health under iron-limiting conditions (Rosconi et al., 2016). These and other features support the potential of *Variovorax* as an agricultural probiotic (Natsagdorj et al., 2019; Bessadok et al., 2020).

We demonstrated that the *Variovorax* was not only capable of aiding the growth of a thiamine-auxotrophic microorganism but was able to produce extracellular concentrations of thiamine up to 100 mg/L. Bacteria typically require only 0.1-10 ng/ml for optimal growth (Newman et al., 1985; Kouno et al., 1973) and natural and most engineered overproducers typically produce only 1-2 mg/L of extracellular thiamine (Bali et al., 2020, Tokui et al., 2011; Schyns et al., 2005). However, thiamine deregulatory mutants have been developed for biotechnological application (Schyns et al., 2005; Goese et al., 2012) that can produce up to 300 mg/L of thiamine. These mutations involve the deletion of the thiamine-monophosphate kinase (*thiL*) responsible for the phosphorylation of thiamine-phosphate, a thiamine permease (*yuaJ*) and active transporter responsible specifically for the influx of thiamine (*ykoD*). Analysis of the isolated *Variovorax* thiamine-monophosphate kinase (*thiL*) protein sequence appears functional based on a comparison with the crystal structure and sequence analysis performed by McCulloch et al. (2008). The active site residues of our *Variovorax’s thiL* appear conserved and the additional conserved residues identified by McCulloch and co-authors were largely maintained. This supports that the regulation of this operon is functional. We suggest that *Variovorax* has evolved to produce large quantities of thiamine to support dependent organisms some of which, in turn, produce pantothenate.

The growth of *Variovorax* in minimal media could be stimulated through the addition of pantothenate. Supplementing greater concentrations of pantothenate led to a shorter duration of lag phase with higher supported cell densities. Thiamine overproduction was then observed during this lag phase. The observed change from thiamine production to consumption by *Variovorax*, at the start of exponential growth, confirms that the thiamine biosynthesis operon can be properly regulated and is not “constitutively on” as occurs in dysregulated mutants (Goese et al., 2012). We hypothesize that the production of high concentrations of thiamine during lag phase may have evolved in this *Variovorax* to stimulate the growth of primarily pantothenate-producing bacteria. *Variovorax* is correlated with two pantothenate producers. Although *Crypotoccus* did not occur frequently enough to be included in the network, it probably exemplifies this co-dependency. The extreme thiamine-producing phenotype and the prevalence of this operon across the *Variovorax* genus suggest the potential of *Variovorax* for biotechnological vitamin production.

Our integration of network analysis with pathway reconstruction from genome-resolved metagenomics represents an easily implementable tool for identifying interacting microbial community members and their mechanism of interaction. The approach demonstrates the value of metagenomic datasets from sample series and is readily implementable in future studies of any microbial consortia.

## Conclusion

Using laboratory consortia experiments and co-cultures of isolated microorganisms, we demonstrate the predictive power of combined network analyses, genome-resolved metagenomics, and pathway reconstruction for the elucidating interactions in complex microbial communities. We provide evidence for interdependencies such as between species of symbiotic *Saccharibacteria*, *Actinobacteria* and *Variovorax*, and for vitamin exchange, such as between *Variovorax* and *Cryptococcus*. Our results predict that *Variovorax* in our consortia has evolved to overproduce thiamine to support organisms that depend on it for this vitamin and provide evidence to suggest that thiamine production is likely an important trait for these bacteria in many ecosystems.

## Materials and Methods

### Bioreactor operation and sampling

The bioreactors used to study this microbial consortium were operated as described by Kantor et al. (2015), Kantor et al. (2017), Rahman et al. (2017) and Huddy et al. (2021). Briefly, 1 L glass stirred-tank reactors were inoculated with homogenised biofilm and planktonic samples taken from long-running thiocyanate-degrading bioreactors at the University of Cape Town. The reactors were stirred using a pitched-blade impeller at 270 rpm and sparged with filtered air at 0.9 L/min. The bioreactors were fed continuously with deionised water containing 0.28 mM KH_2_PO_4_ and were buffered to pH 7.0 using 5N KOH. Variable concentrations of sodium-thiocyanate (50 - 2000 mg/L) were supplied in the medium as the predominant electron donor. Molasses was included in some of the bioreactor mediums at 0.15 g/L while being omitted in others. All of the bioreactors were operated continuously at hydraulic retention times varying from 12 - 42 hours. Biofilm and planktonic samples were recovered from the bioreactors at chemical steady states as described by the aforementioned studies.

### Microbial genomes and annotation

Read assembly, and genome binning were performed as previously described (Kantor et al., 2015; Kantor et al., 2017; Rahman et al., 2017; Huddy et al., 2021) and genomes are available https://ggkbase.berkeley.edu/project_groups/thiocyanate_bioreactor and in NCBI as previously documented. The MAGs from each metagenome were pooled and a final genome set was produced using dRep (Olm et al., 2017) at 95% ANI. Open reading frames (ORFs) were predicted using Prodigal’s (Hyatt et al., 2010) and annotated using Hmmscan (http://hmmer.org/) against TIGRFAM and PFAM databases. Vitamin pathways were recovered from KEGG website and applicable literature (Kanehisa et al., 2000). The capacity to produce a vitamin was determined based on a minimum set of genes representing different stages of each pathway. These minimum genes necessary for the deemed presence of a pathway are discussed in supplementary materials.

### Network analysis

Reads were mapped to the dereplicated genome set using CoverM (https://github.com/wwood/CoverM). Correlations and associated p-values were calculated using FastSpar (Watts et al., 2019) which implements the SparCC algorithm (Friedman & Alm, 2012). Network analysis was performed using correlations greater or equal to 0.35. No negative correlations were considered. The Network analysis was performed using NetworkX (Hagberg et. al., 2008) and was used to calculate the Betweeness-centrality (eq1) of each node and the overall network density (eq 2) (2 * #edges) /(Total nodes * (Total nodes −1).

A third metric was developed in this study for the quantification of the importance of a vitamin producer within the network. This was termed the ‘vitamin-hub metric’ (eq3) and is calculated by multiplying the degree of a node, with the capacity to synthesise a given vitamin, by the number of edges it shares with nodes which are predicted to be auxotrophic for the given vitamin. This product is divided by the mean number of shared edges between each of the node’s neighbours and other nodes which are also predicted to synthesise the vitamin. One is added to the bottom term to prevent the metric from becoming undefined when this mean value is equal to zero. Networks were visualised using Cytoscape v3.8.2 (http://www.cytoscape.org/)

### GTDB Variovorax annotation and phylogenetic analyses

Species representatives in GTDB (r207_v2; https://gtdb.ecogenomic.org/searches?s=al&q=Variovorax) were annotated as described above using KEGG HMMs and their capacity to synthesise thiamine evaluated. Genomes classified as Variovorax were then dereplicated at 98% average nucleotide identity (~subspecies level) using dRep (Olm et al., 2017). GToTree (Lee, 2019) was used to identify, align, trim and concatenate 16 universal ribosomal proteins from the Variovorax genomes. A maximum-likelihood Phylogenetic reconstruction of these alignments was performed using IQtree (-st AA -nt 48 -bb 1000 -m LG+G4+FO+I”) (Nguyen et al., 2015). The phylogenetic tree was then annotated in iTol (v 6.5.8; Letunic and Bork, 2007). The contamination of these genomes was estimated using CheckM v1.2.1 (Parks et al., 2015). Complete thiamine biosynthesis pathways contained all Kegg annotations for (i) thiazole production, (ii) pyrimidine synthesis, (iii) the linking of these two moieties and (iv) the kinases involved in the overall biosynthesis process. Near-complete pathways were those that only lacked a phosphomethylpyrimidine synthase (*thiC*, K03147) from this pathway.

### Microbial isolation, sequencing and assembly

Microbial isolates were recovered from an in-house thiocyanate-degrading bioreactor by serial dilution in sterile 1x PBS buffer and spread plating onto R2A solid agar medium. Recovered isolates were grown in liquid R2A medium and 600 uL combined at 1:1 with 50% (v/v) glycerol before being stored at −80 °C. The remaining cell culture was pelleted at 10 000 g for 10 minutes and used for subsequent DNA extraction. Total DNA was extracted from the isolates using a DNAeasy^®^ PowerSoil^®^ Pro DNA extraction kit (Machery-Nagel, Germany). Paired-end Illumina TruSeq libraries with fragment sizes of 500 bp were sequenced on an Illumina NovaSeq to yield 250 bp paired-end reads. Read assembly and genome annotation were conducted as described above.

### Isolate culturing and co-culturing

The recovered isolates were screened for thiamine auxotrophies using solid M9 minimal media (0.4 g/L glucose, 2 mM MgSO_4_, 0.1 mM NaCl, 0.048 mM Na_2_HPO_4_.7H_2_O, 0.022 mM KH_2_PO_4_, 0.017 mM NaCl, 15 g/L agar), with and without the addition of 0.5 mg/L thiamine. The *Cryptococcus* species identified as a possible thiamine auxotroph was grown in a 96-well plate in liquid M9 minimal media with (n=6) and without (n=6) the addition of 0.5 mg/L thiamine. Growth was quantified spectrophotometrically at 600 nm.

To evaluate the effect of the growth of Variovorax on *Cryptococcus*, and vice versa, each of these isolates were grown in pure culture and in co-culture in minimal media. Initially, the isolates were grown in R2A medium overnight, spun down at 3 000 g for 3 minutes and washed in M9 minimal media before being inoculated into 10 ml M9 minimal media each at an initial OD_600_ of 0.035. The pure and co-cultures were allowed to grow for 48 hours before passaging into fresh M9 media, and this repeated for a total of 4 passages. Immediately following each passage, viable cell densities were determined by serial dilution in M9 media and spread plating onto R2A media agar. These plates were incubated at 30 °C for 48 hours and colonies were counted and differentiated based on colony morphology (*Cryptococcus* white, *Variovorax* yellow).

Variovorax was cultured in M9 minimal media in the presence of varying concentrations of pantothenate using the methodology described above but allowed to grow for a total of 9 days. The cultures were performed with four replicates and were sampled aseptically every 24 hours and quantified for cell optical density. Subsequently, cells were pelleted at 10 000 g for 3 minutes in a benchtop centrifuge and filter sterilised using a 0.22 um filter. These solutions were then used for the quantification of extracellular thiamine.

### Thiamine and Pantothenic acid quantification

The quantity of thiamine produced by Variovorax was quantified during the growth in 75% fresh minimal media and 25% filter-sterilized spent medium from the final passage of the co-culture experiment as well as during the growth of Variovorax in M9 minimal media with varying concentrations of pantothenate. Thiamine was quantified on an Agilent 1200 liquid chromatograph equipped with a Agilent ZORBAX Eclipse Plus C-18 column with a 5 um particle size, maintained at 35 °C. Using a two phase mobile phase of (A) 25 mM NaH_2_PO_4_ (pH = 2.5), and (B) pure methanol. The flow rate was 1.0 mL/min. The mixture of these mobile phases were as follows: 0% mobile phase B from time of injection, 50% mobile phase B from 1.0 minute to 75% mobile phase B from 10 to 25 minutes. We identified vitamins using a G1362A Refractive Index Detector at 220 nm. Vitamin standards were obtained from Sigma-Aldrich^®^.

## Acknowledgements

This material by m-CAFEs Microbial Community Analysis and Functional Evaluation in Soils, a Science Focus Area led by Lawrence Berkeley National Laboratory is based upon work supported by the U.S. Department of Energy, Office of Science, Office of Biological and Environmental Research under contract number DE-AC02-05CH11231. We thank Yi Liu for assistance with HPLC and Dr Maria Lukarska for assistance with *thiL* sequence comparison.

## Supplementary figures

**Fig S1.**
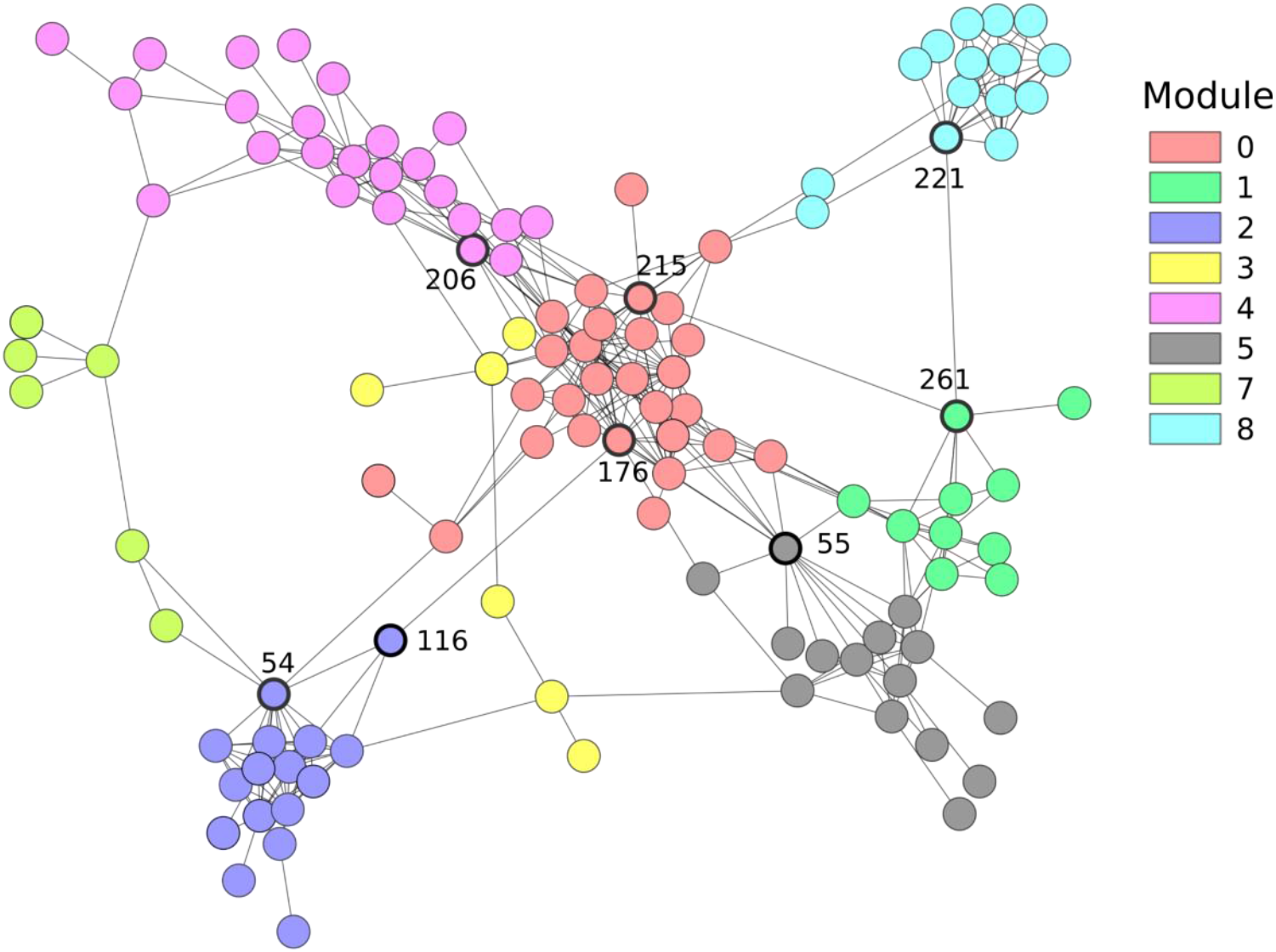
Network analysis based on sparCC correlation across 92 metagenomes, as shown in Figure 2A, showing modules in differing colors and delineated network hub, based on elevated betweeness centrality, with bold node borders. See Tabls S2 for further network and node information.

**Figure S2.**
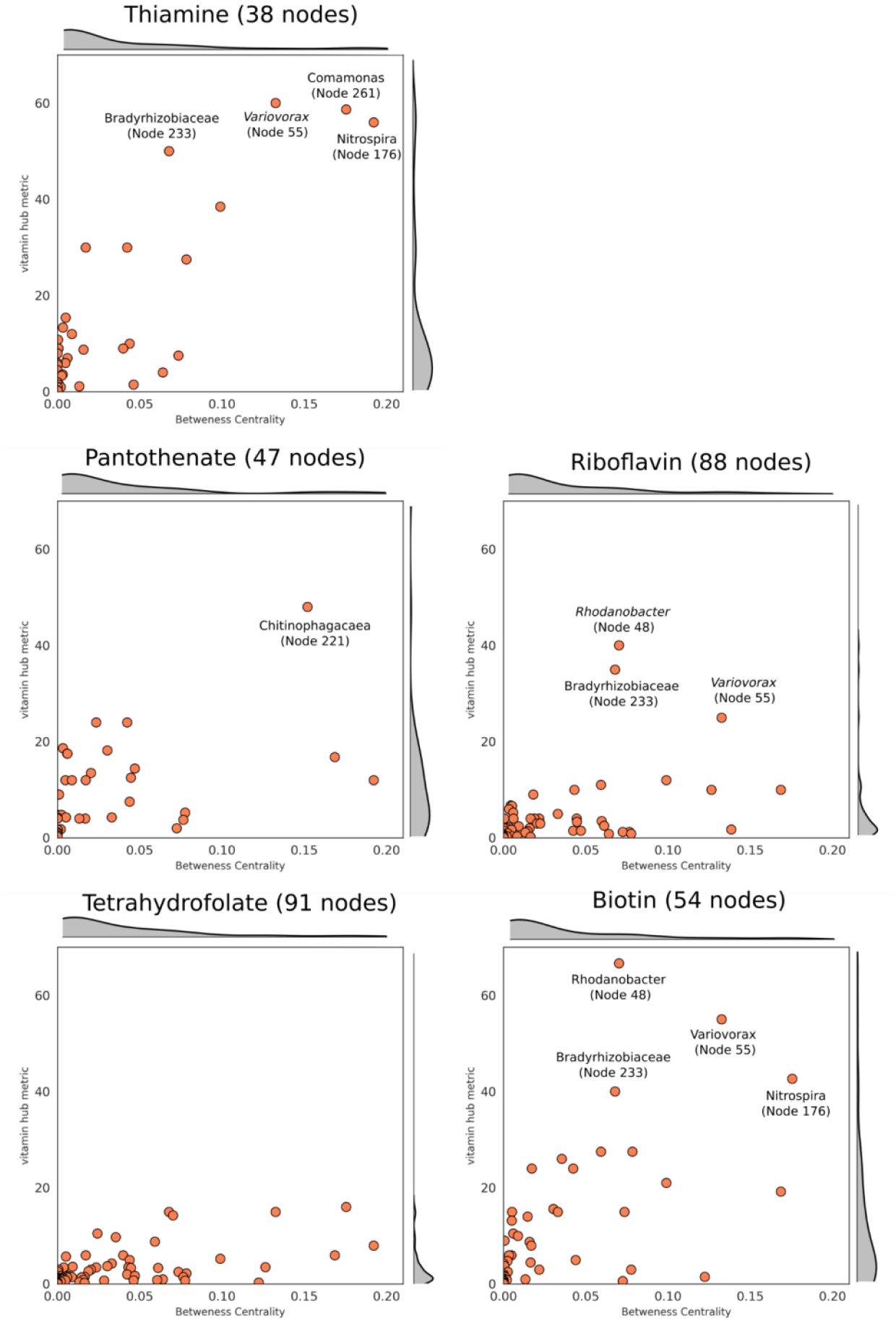
Vitamin hub metric of thiamine, pantothenate, riboflavin, tetrahydrofolate and biotin based on the observed distribution of vitamin producers across the network shown in Figure 2A.

**Fig S3.**
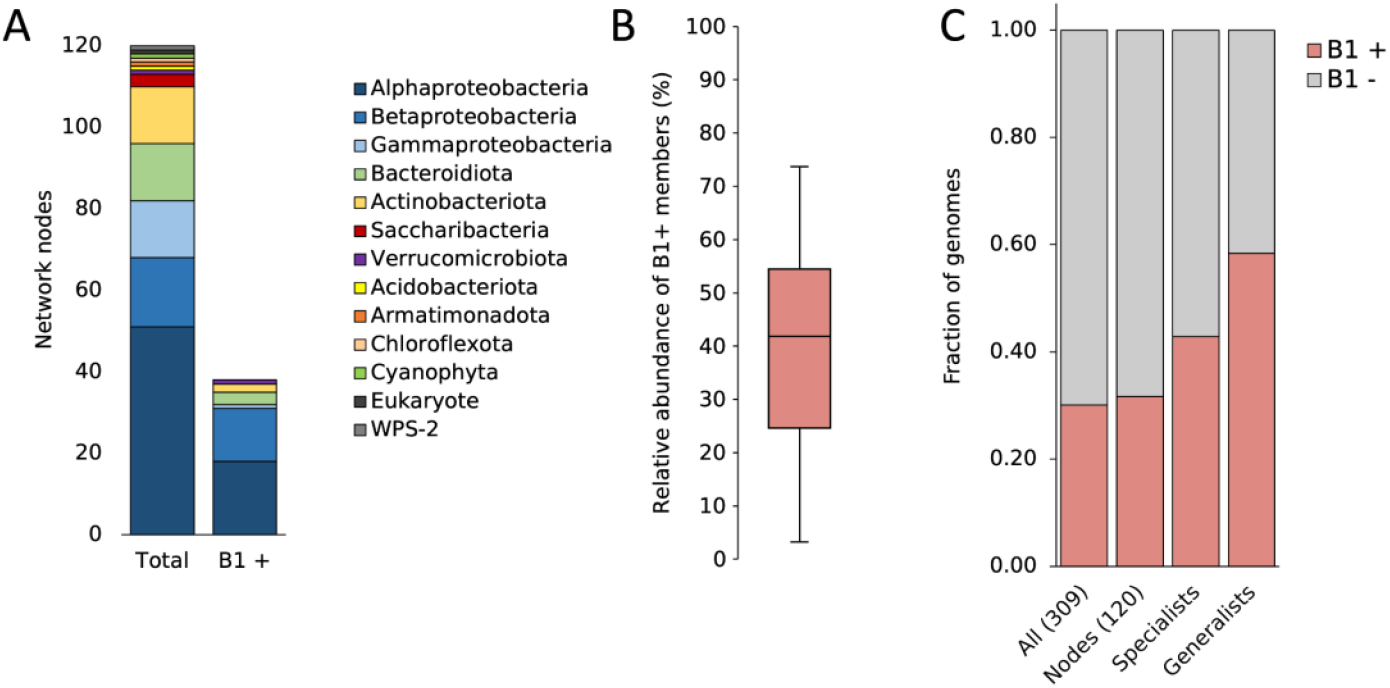
(A) The taxonomy of the organisms represented in the correlation network and those in the network capable of thiamine biosynthesis. (B) The distribution in the proportion which encodes their own thiamine biosynthesis across the 92 microbial communities. (C) The proportion of all 309 bacteria, the 120 bacteria represented in the network and of the specialists and generalists which endcode their own B1. Generalists are defined as organisms which are found in 75% of the 92 metagenomes and have a mean relative abundance of greater than 1%. Specialists are defined as those which are present in 25% or fewer of the metagenomes but have a mean relative abundance of greater than 1%.

**Figure S4.**
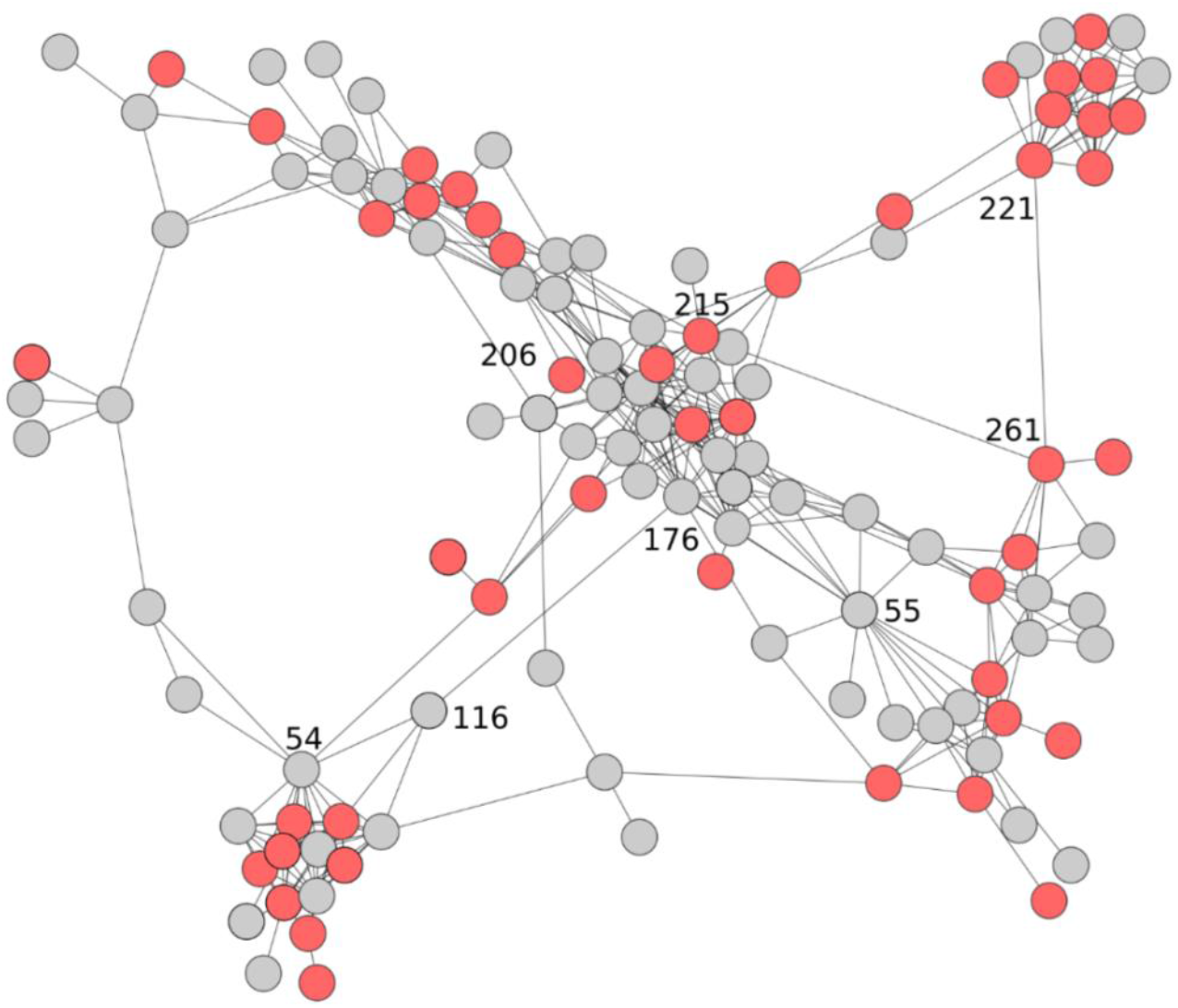
Network analysis based on sparCC correlation across 92 metagenomes, as shown in Figure 2A, showing organisms which encode for the biosynthesis of pantothenic acid in red and absent in grey.

**Figure S5.**
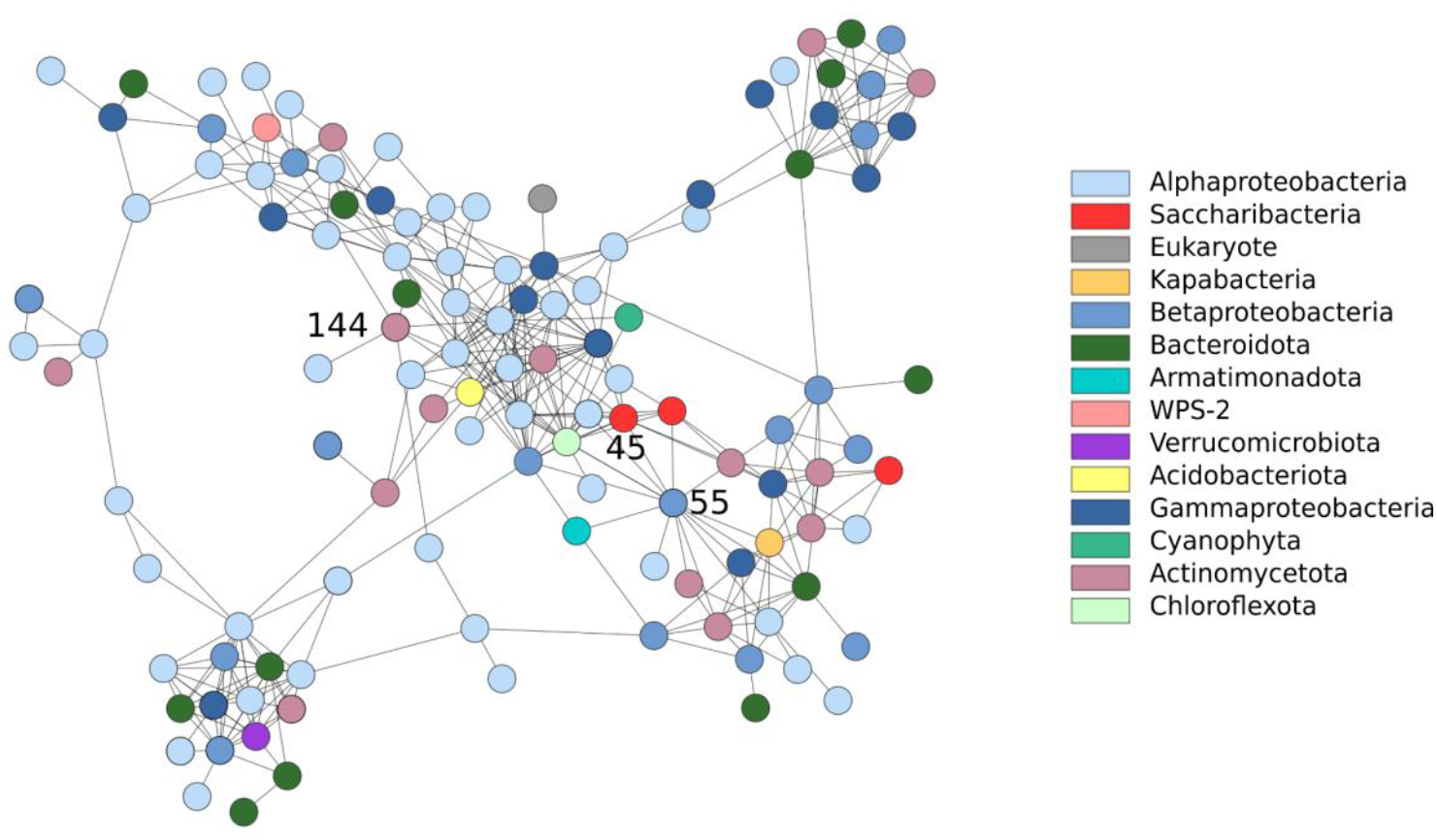
Figure 5 with greater node separation: *Variovorax* (Node 55) is correlated with a *Saccharibacteria* (Node 45) as illustrated in the network plot showing phyla annotations. *Microbacterium* (Node 144) and *Saccharibacteria* both have multiple genes which encode proteins which require thiamine as a cofactor - neither of these genomes encodes for thiamine biosynthesis

